# VIBRIO CHOLERAE ADAPTS TO SESSILE AND MOTILE LIFESTYLES BY CYCLIC DI-GMP REGULATION OF CELL SHAPE

**DOI:** 10.1101/2020.04.15.043257

**Authors:** Nicolas L. Fernandez, Nguyen T. Q. Nhu, Brian Y. Hsueh, Joshua L. Franklin, Yann S. Dufour, Christopher M. Waters

## Abstract

The cell morphology of rod-shaped bacteria is determined by the rigid net of peptidoglycan forming the cell wall. While *V. cholerae* grows into a curved shape under most conditions, straight rods have been observed. However, the signals and regulatory pathways controlling cell shape transitions in *V. cholerae* and the benefits of switching between rod and curved shape have not been determined. We demonstrate that cell shape in *V. cholerae* is regulated by the bacterial second messenger cyclic dimeric guanosine monophosphate (c-di-GMP) by repressing expression of *crvA*, a gene encoding an intermediate filament-like protein necessary for curvature formation in *V. cholerae.* This regulation is mediated by the transcriptional cascade that also induces production of biofilm matrix components, indicating that cell shape is coregulated with *V. cholerae*’s induction of sessility. Wild-type *V. cholerae* cells adhering to a surface lose their characteristic curved shape to become as straight as cells lacking *crvA* while genetically engineering cells to maintain high curvature reduced microcolony formation and biofilm density. Conversely, straight *V. cholerae* mutants have reduced speed when swimming using flagellar motility in liquid. Our results demonstrate regulation of cell shape in bacteria is a mechanism to increase fitness in planktonic or biofilm lifestyles.

## Introduction

Morphology is an important feature for every form of life as it dictates how organisms interact with their physical world. Various aspects of morphology such as shape, length, and the presence of appendages are subject to selective pressures and contribute to adaptation to specific ecological niches (1). Not surprisingly, bacteria take on diverse shapes from simple rods and cocci to helices and curves; however, the fitness benefits of each shape is not always well understood (2).

Shape in bacteria is largely determined by the structure of the peptidoglycan (PG) cell wall as bacteria without PG adopt pleiomorphic, or L-form, shapes (3, 4). A predominant morphology in Gram-negative bacteria is the rod shape. During growth of a rod-shaped cell, new PG subunits are added to the growing PG layer by continuous rounds of PG cleavage and subunit insertion at the mid-cell (3). The curved-rod shape is common in many aquatic organisms, such as the fresh-water bacterium *Caulobacter crescentus* and the opportunistic human pathogen *Vibrio cholerae*. In his 1928 book titled “Morphologic Variation of the Cholera Vibrio”, Arthur Henrici measured *V. cholerae* shape as a function of time during a typical bacterial growth curve and observed shapes ranging from spheres to straight and curved rods (5). Research by Bartlett and co-workers identified that the gene *crvA* encoding the periplasmic intermediate filament-like protein CrvA introduces curvature by altering PG insertion rates (6, 7). Mutants lacking *crvA* grow as straight rods, have reduced migration in soft agar, and are less virulent in animal models of infection (7). The adjacent open reading frame to *crvA,* annotated as *crvB,* functions with CrvA and, together, CrvAB are sufficient to produce curvature in normally straight cells (8). This work and others suggest bacterial cell shape determination may be a regulated process, yet what regulatory pathways are used to change shape and potential ecological benefits of such changes are not understood (5, 7).

In fresh and salt-water environments, *V. cholerae* transitions between a motile state powered by a single flagellum or a sessile biofilm lifestyle attached to chitin-coated plankton in dense communities of bacteria encased in a matrix of exopolymeric substances (EPS) (9). In the majority of bacteria, flagellar motility and biofilm formation is regulated by cyclic dimeric guanosine monophosphate (c-di-GMP) signaling (10). In *V. cholerae,* c-di-GMP produced by enzymes called diguanylate cyclases (DGCs) binds to and activates the transcription factors VpsR and VpsT (11, 12). These transcription factors then induce expression of the operons encoding the enzymes necessary for synthesizing *Vibrio* polysaccharide (VPS), the major EPS component of the biofilm matrix, and matrix-associated proteins resulting in mature biofilm formation (13–16). In addition, c-di-GMP represses motility by inhibiting transcription of the flagellar biosynthesis genes, repressing the expression of the transcription factor TfoY by binding to a c-di-GMP-dependent riboswitch, and inducing MshA pilus extension mediating surface attachment (17, 18).

In this study, we report that elevated intracellular c-di-GMP concentrations straighten *V. cholerae* by decreasing CrvA expression. Locking *V. cholerae* in a curved morphology reduced the formation of microcolonies and biofilm formation. Conversely, *V. cholerae* cells locked in a straight morphology had reduced swimming speed when compared to the curved wild-type cells. Therefore, curved and straight cell morphologies are optimal for motility and biofilm formation, respectively, and our results demonstrate how bacteria can actively control cell shape to adapt to different lifestyles.

## Results

### High Intracellular C-di-GMP Reduces Cell Curvature

When exploring the impact of c-di-GMP on stress response pathways in *V. cholerae,* we noted that strains with high intracellular concentrations of c-di-GMP were more frequently straight rods (19, 20). We explored this finding by quantifying cell shape parameters under different intracellular c-di-GMP concentrations utilizing strains that ectopically express an IPTG inducible diguanylate cyclase (DGC) (QrgB) that synthesizes c-di-GMP or a catalytically inactive DGC (QrgB*). We have previously shown such expression of QrgB in *V. cholerae* produces physiologically relevant concentrations of c-di-GMP (18). C-di-GMP did not measurably alter cell length or width (Extended Data Figure 1). However, the curvature of the medial axis of cells decreased in a strain harboring the active DGC when no IPTG was added, due to leaky expression of the inducible promoter, (p-val = 5.6e-3), and an even further reduction of curvature was observed when IPTG was added to increase QrgB expression (p-val = 4.3e-2), suggesting that high c-di-GMP concentrations cause cells to straighten (Figure 1).

**Figure 1:**
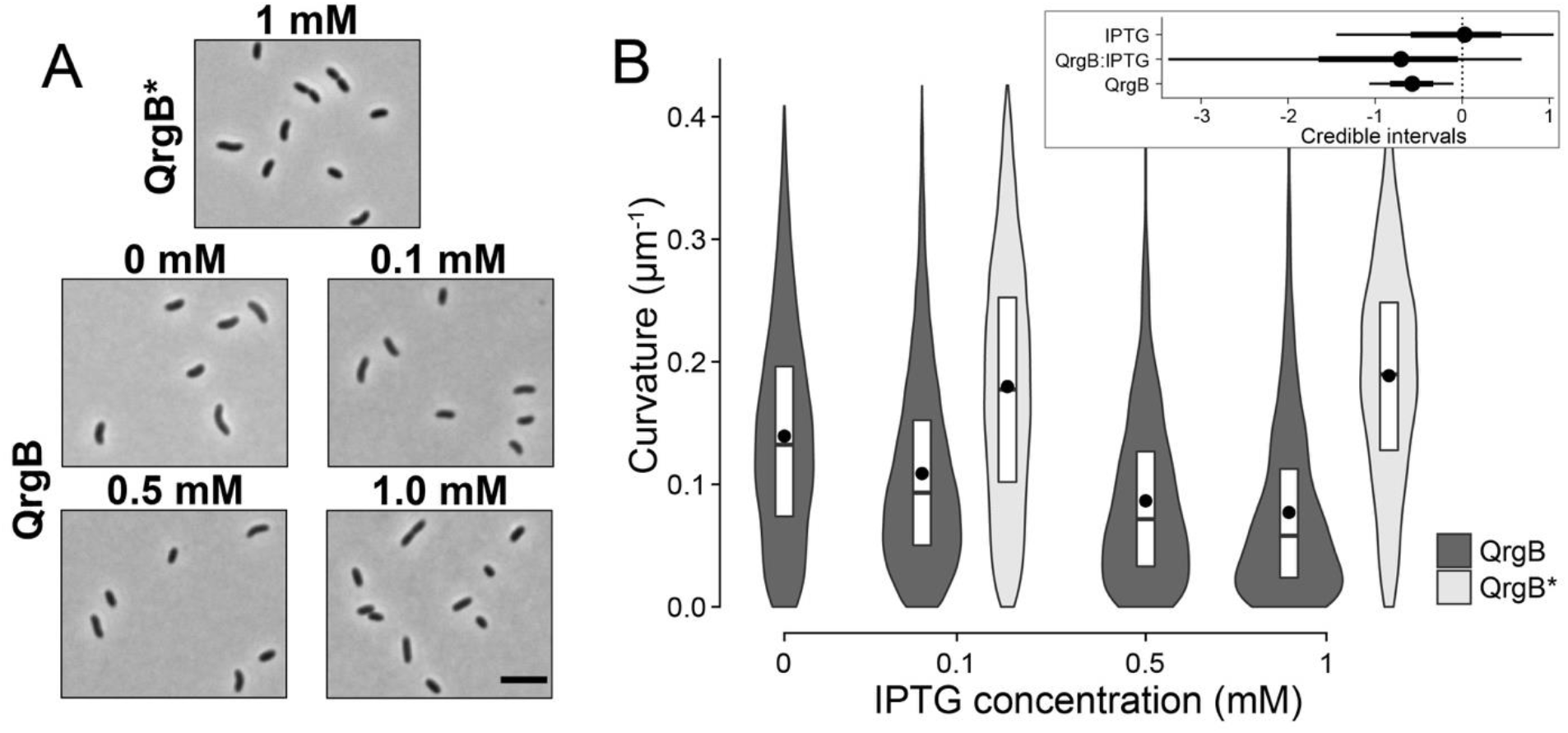
C-di-GMP Decreases Cell Curvature in a Dose-Dependent Manner. A) Representative phase-contrast micrographs of early stationary phase cells harboring a plasmid encoding an IPTG-inducible inactive DGC (QrgB*) with 1 mM IPTG and an IPTG-inducible active DGC (QrgB) with 0, 0.1, 0.5, and 1 mM IPTG. The bar on the last image of the panel is 5 μm. B) Distributions of cell curvature as a function of IPTG concentration in populations expressing inactive DGC (QrgB*, light) or active DGC (QrgB, dark). The box represents the first, second, and third quartiles. The dot represents the mean. Each distribution represents between 1,000 and 1,200 cells analyzed and pooled from two to three separate experiments. Insert) Credible intervals of the contribution of each experimental factor to changes in the medians (dot = mean, thick line = 90% CI, thin line = 98% CI).

**Extended Data Figure 1:**
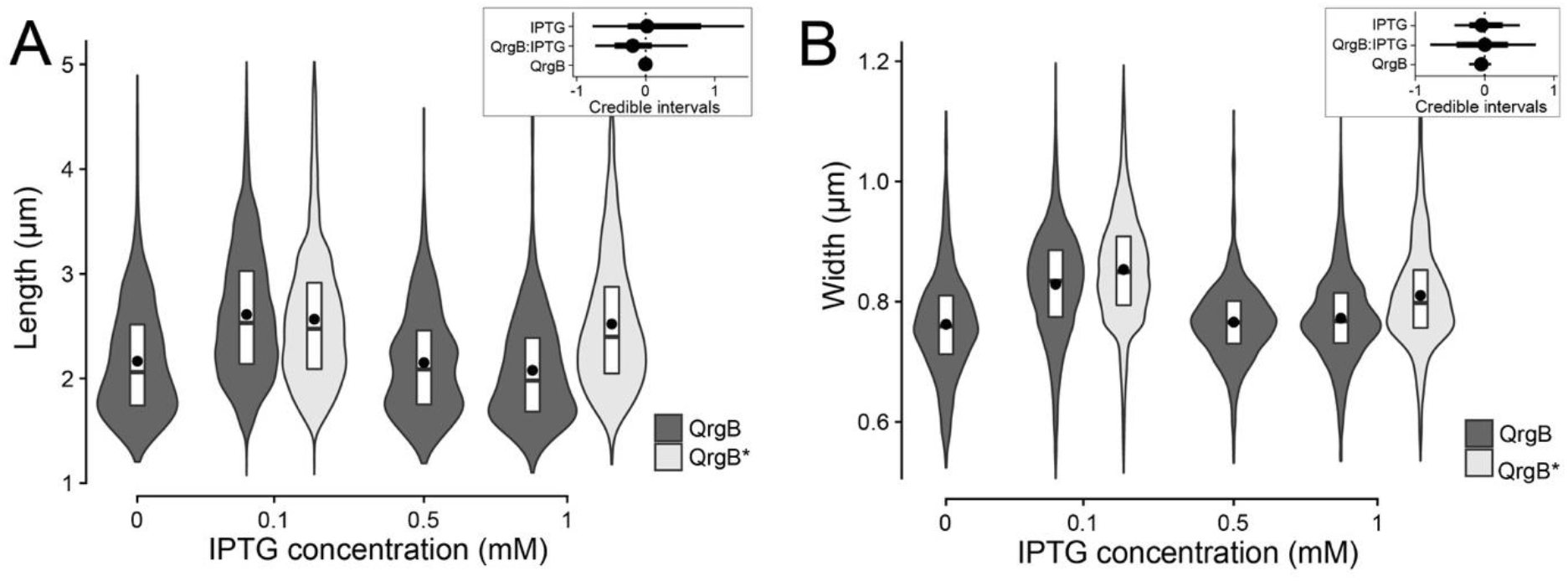
C-di-GMP Does not Significantly Alter Cell Length or Width. Distributions of cell length **(A)** and width **(B)** as a function of IPTG concentration in populations expressing inactive DGC (QrgB*, light) or active DGC (QrgB, dark). The box represents the first, second, and third quartiles. The dot represents the mean. Each distribution represents between 1,000 and 1,200 cells analyzed and pooled from two to three separate experiments. **Insert**) Credible intervals of the contribution of each experimental factor to changes in the medians (dot = mean, thick line = 90% CI, thin line = 98% CI).

### The C-di-GMP-Dependent Transcription Factors, VpsT and VpsR, Control C-di-GMP Mediated Changes to Cell Curvature

In *V. cholerae*, three c-di-GMP dependent transcription factors (VpsR, VpsT, and FlrA) and two c-di-GMP-dependent riboswitches (Vc1 and Vc2), are known to regulate genes that elicit diverse phenotypes such as biofilm formation, motility, DNA repair, and catalase production (21, 14, 22, 11, 17, 23, 20, 24, 19). We hypothesized that VpsT or VpsR were involved in the c-di-GMP-dependent regulation of cell curvature because of their roles in regulating multiple c-di-GMP-dependent phenotypes (19, 20, 25). To test this hypothesis, we measured curvature in the parent strain and isogenic *ΔvpsR* mutant at high (QrgB) and unaltered (QrgB*) concentrations of c-di-GMP. The c-di-GMP mediated decrease in curvature was lost in the *ΔvpsR* mutant (Figure 2). Expression of VpsR from a multicopy plasmid in the *ΔvpsR* mutant restored c-di-GMP-dependent inhibition of cell curvature (p-val = 2.6e-2). Similarly, c-di-GMP inhibition of cell curvature was lost in the *ΔvpsT* mutant (Figure 2). In this case, expression of VpsT from a multicopy plasmid resulted in decreased cell curvature at both high and unaltered c-di-GMP concentrations (p-val = 4.3e-3) while retaining some sensitivity to c-di-GMP (pval = 5.7e-3). VpsT is a c-di-GMP dependent transcription factor, but this result suggests that high levels of VpsT expression are sufficient to decrease curvature at the unaltered concentrations of c-di-GMP (11). As VpsR directly activates *vpsT* transcription, we hypothesized the role of VpsR in controlling cell shape is to increase VpsT expression, which reduces curvature (12). We tested this hypothesis by expressing VpsT in the *ΔvpsR* mutant and found that cells appeared straight regardless of intracellular c-di-GMP concentrations (Figure 2). Moreover, induction of VpsR in the *vpsT* mutant did not restore c-di-GMP inhibition of curvature. Collectively, these data indicate that at high c-di-GMP conditions, VpsR inhibits curvature indirectly by inducing transcription of *vpsT* while VpsT is sufficient for inhibiting curvature.

**Figure 2:**
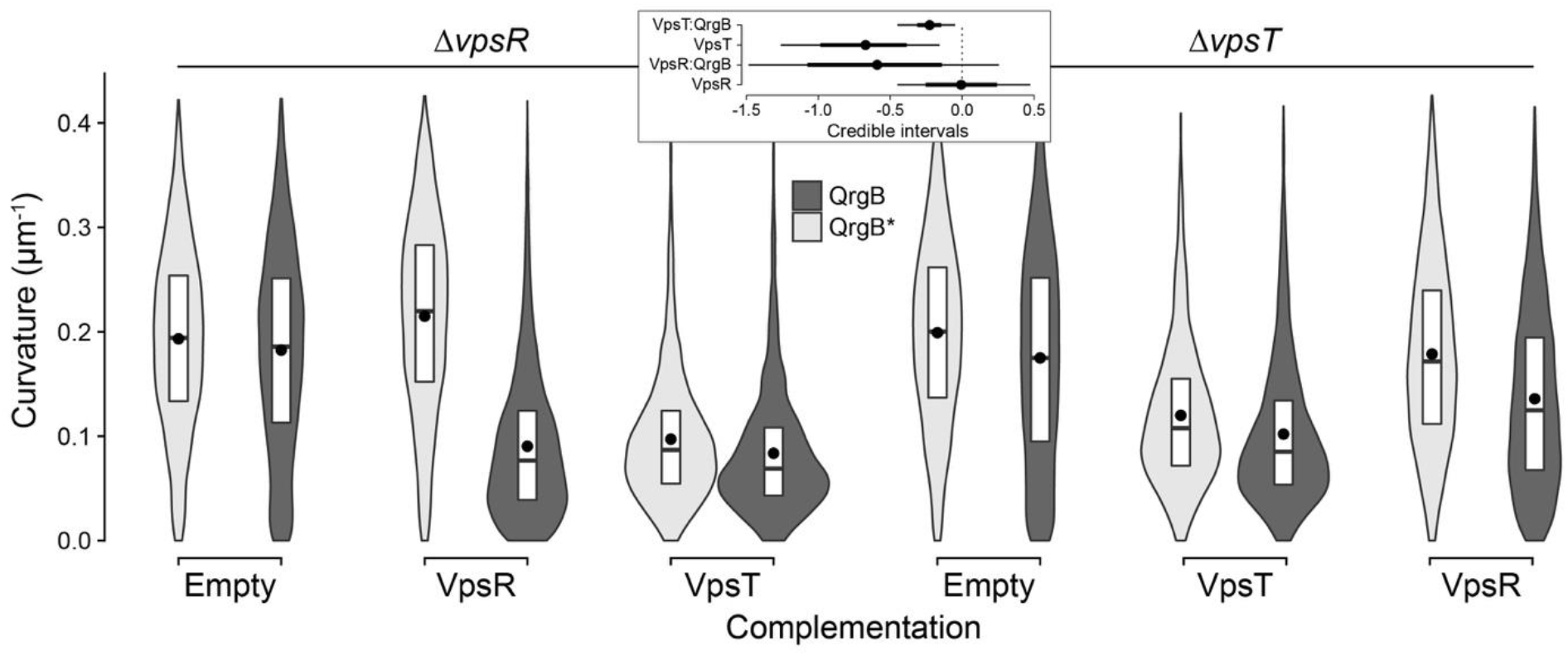
The Transcription Factors VpsR and VpsT are Necessary for C-di-GMP Dependent Loss of Curvature. Distributions of cell curvature from different mutant strains in populations expressing inactive DGC (QrgB*, light) or active DGC (QrgB, dark). The *ΔvpsR* strain was complemented with either an empty plasmid or a plasmid expressing VpsR or VpsT. The *ΔvpsT* strain was complemented with either an empty plasmid or a plasmid expressing VpsR or VpsT. The box represents the first, second, and third quartiles. The dot represents the mean. Each distribution represents between 1,000 and 1,200 cells analyzed and pooled from two to three separate experiments. **Insert**) Credible intervals of the contribution of each experimental factor to changes in the medians (dot = mean, thick line = 90% CI, thin line = 98% CI).

### C-di-GMP Decreases CrvA Expression to Inhibit Curvature

In *V. cholerae,* cell curvature is generated by the intermediate-filament like protein CrvA by decreasing net growth on the minor axis relative to the major axis (7). We hypothesized that c-di-GMP-dependent inhibition of cell curvature was due to negative regulation of *crvA* transcription. To test this hypothesis, we generated a transcriptional reporter of the *crvA* promoter (P_*crvA*_, 358bp upstream of the *crvA* transcriptional start site) fused to luciferase. This promoter region was sufficient to complement a *ΔcrvA* mutant when integrated with the *crvA* ORF at a heterologous site on the chromosome (Extended Data Figure 2).

**Extended Data Figure 2:**
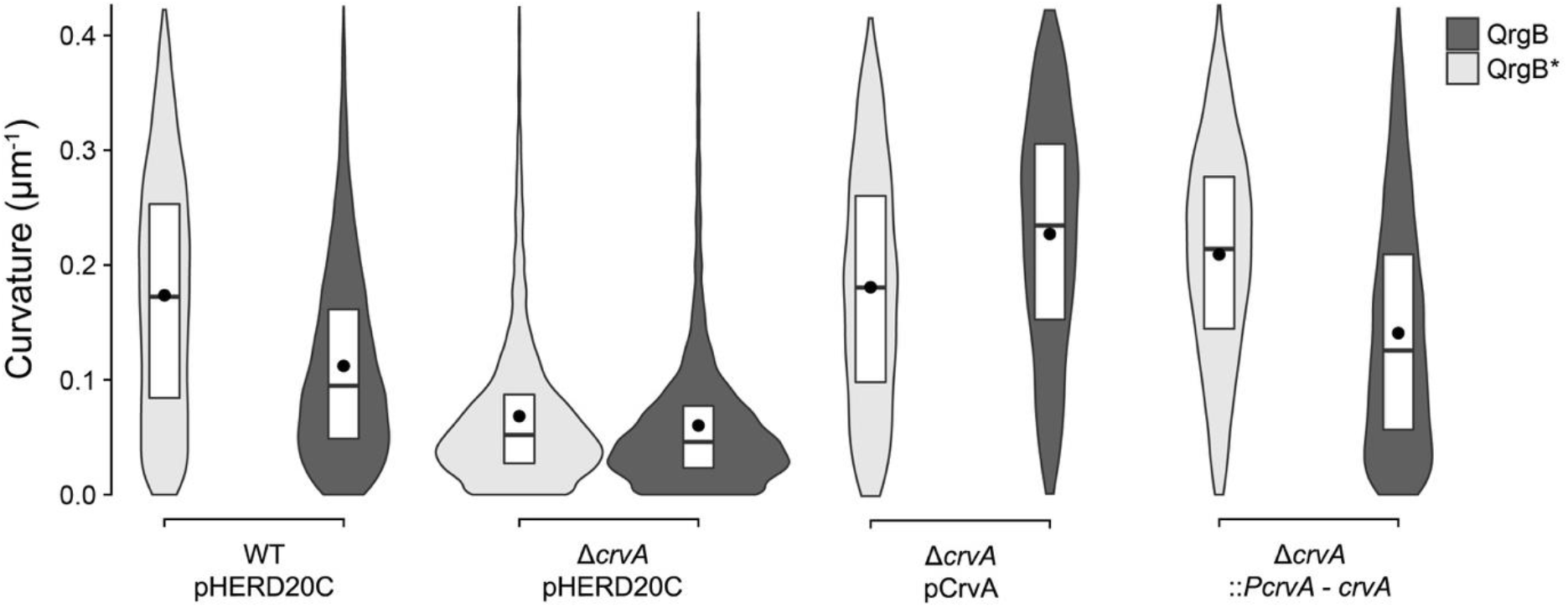
Complementation of *ΔcrvA* Restores Curvature. Distributions of cell curvature from different mutant strains in populations expressing inactive DGC (QrgB*, light) or active DGC (QrgB, dark). The box represents the first, second, and third quartiles. The dot represents the mean. Each distribution represents between 2,000 and 3,000 cells analyzed and pooled from two to three separate experiments.

Contrary to our hypothesis, *P*_*crvA*_ transcriptional activity was not affected when c-di-GMP concentration was increased through QrgB expression at all culture densities examined (Figure 3A) (difference in the slopes, 95% CI = [−0.11, 0.10]). In agreement with previous studies that observed increased curvature at high cell density (5, 7), *P*_*crvA*_ transcriptional activity increased with cell density (Figure 3A) (p-val < 2.5e-4).

**Figure 3:**
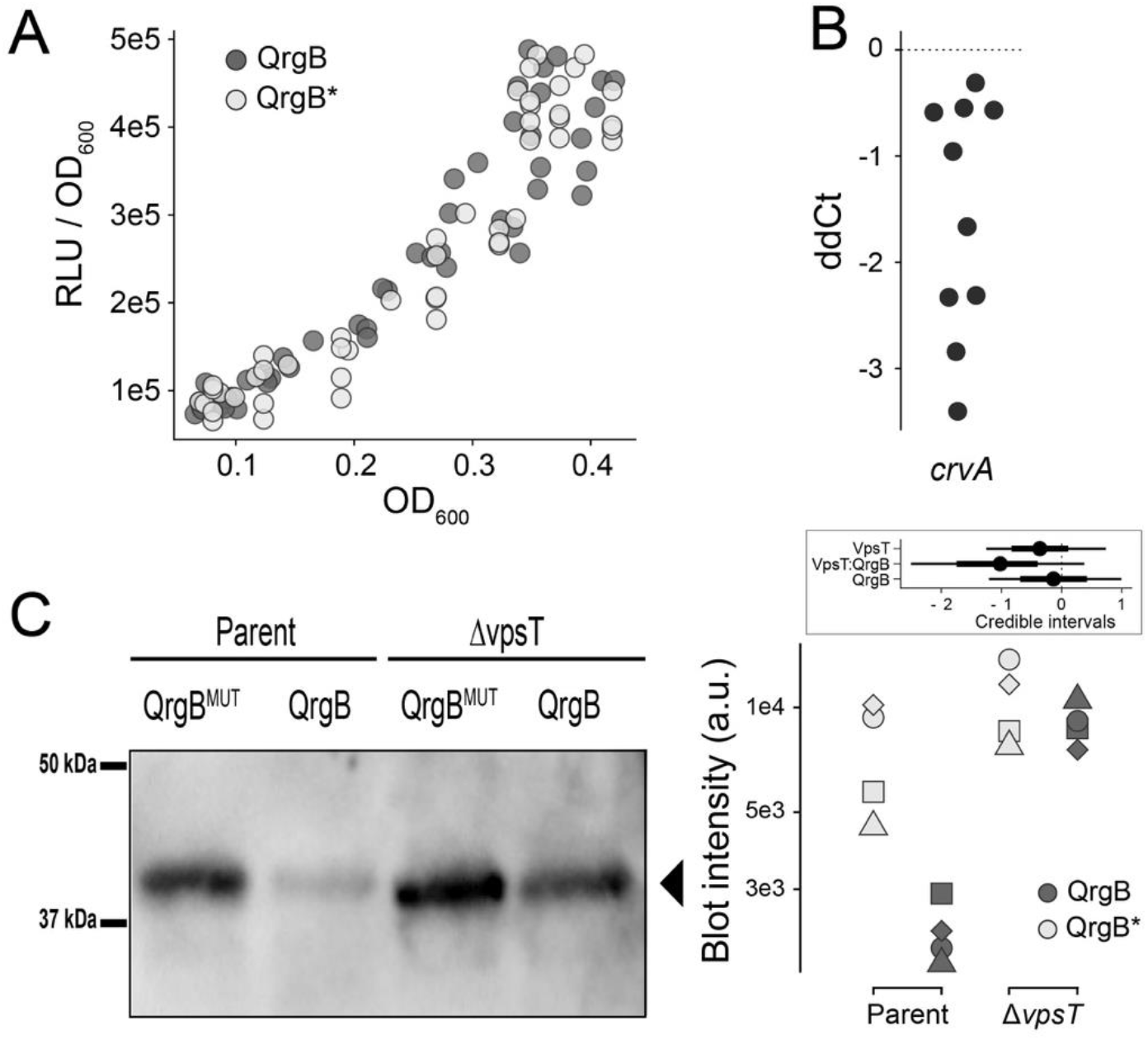
Regulation of CrvA Expression Occurs at the Post-Transcriptional Level. A) Normalized luciferase reporter activity (relative light units) of *P*_*crvA*_ - luciferase transcriptional fusion as a function of cell density (OD_600_) under high (QrgB, dark) and unaltered (QrgB*, light) c-di-GMP conditions. Dots represent pooled data from 4 biological replicates. B) qRT-PCR analysis of relative *crvA* transcript levels between high (QrgB) and unaltered (QrgB*) c-di-GMP conditions. Each dot represents 1 independent replicate. C) Western blot analysis of CrvA-HIS under high (QrgB, dark) or unaltered (QrgB*, light) c-di-GMP conditions in the parent and *ΔvpsT* strains. A representative blot is shown on the left and quantification of band intensities for all replicates on the right. Shapes represents independent replicates. **Insert**) Credible intervals of the contribution of each experimental factor to changes in the medians (dot = mean, thick line = 90% CI, thin line = 98% CI).

Next, we tested if c-di-GMP impacted CrvA accumulation via a post-transcriptional mechanism by quantifying the effects of c-di-GMP on *crvA* mRNA abundance. *crvA* mRNA was ~1.5-fold less abundant high c-di-GMP conditions (Figure 3B) (95% CI = [−2.4, −0.7]). We also quantified the abundance of CrvA fused to a 6-histidine-tag and found a ~3-fold decrease of CrvA protein at high c-di-GMP in the wild-type background (95% CI = [−10, −1.2], but not in the Δ*vpsT* background (95% CI = [−2.4 −0.51]) (Figure 3C). These data demonstrate high c-di-GMP concentrations decreases *crvA* mRNA and CrvA protein levels in a VpsT-dependent manner without altering *crvA* promoter activity, indicating that c-di-GMP negatively controls cellular c curvature by decreasing CrvA abundance via a post-transcriptional mechanism.

### Cell Shape Influences Biofilm Formation at the Single-Cell and Population Level

The ability of *V. cholerae* to attach to surfaces and initiate biofilm formation is dependent on intracellular c-di-GMP and the transcription factors VpsR and VpsT (14, 21, 26). This suggests that c-di-GMP dependent changes to cell shape occurs as *V. cholerae* is initiating biofilm formation concomitantly with VpsT inducing production of VPS and associated biofilm matrix proteins (11, 27, 28). We therefore hypothesized that straight cells of *V. cholerae* would be more adept at forming biofilms. To test this hypothesis, we first grew wild-type *V. cholerae* under static conditions and measured the morphology of attached cells over time starting from initial contact to the early stages of microcolony formation. We chose these timepoints because at later times *V. cholerae* grows in dense clusters with some cells in the biofilm becoming vertical relative to the surface at later stages of biofilm development, making it difficult to segment and quantify curvature from two-dimensional images (29, 30). We observed that over the course of the experiment (7 hours) cells attached to the glass surface became straighter than cells from the initial inoculum (Figure 4) (p-val = 2.3e-2).

**Figure 4:**
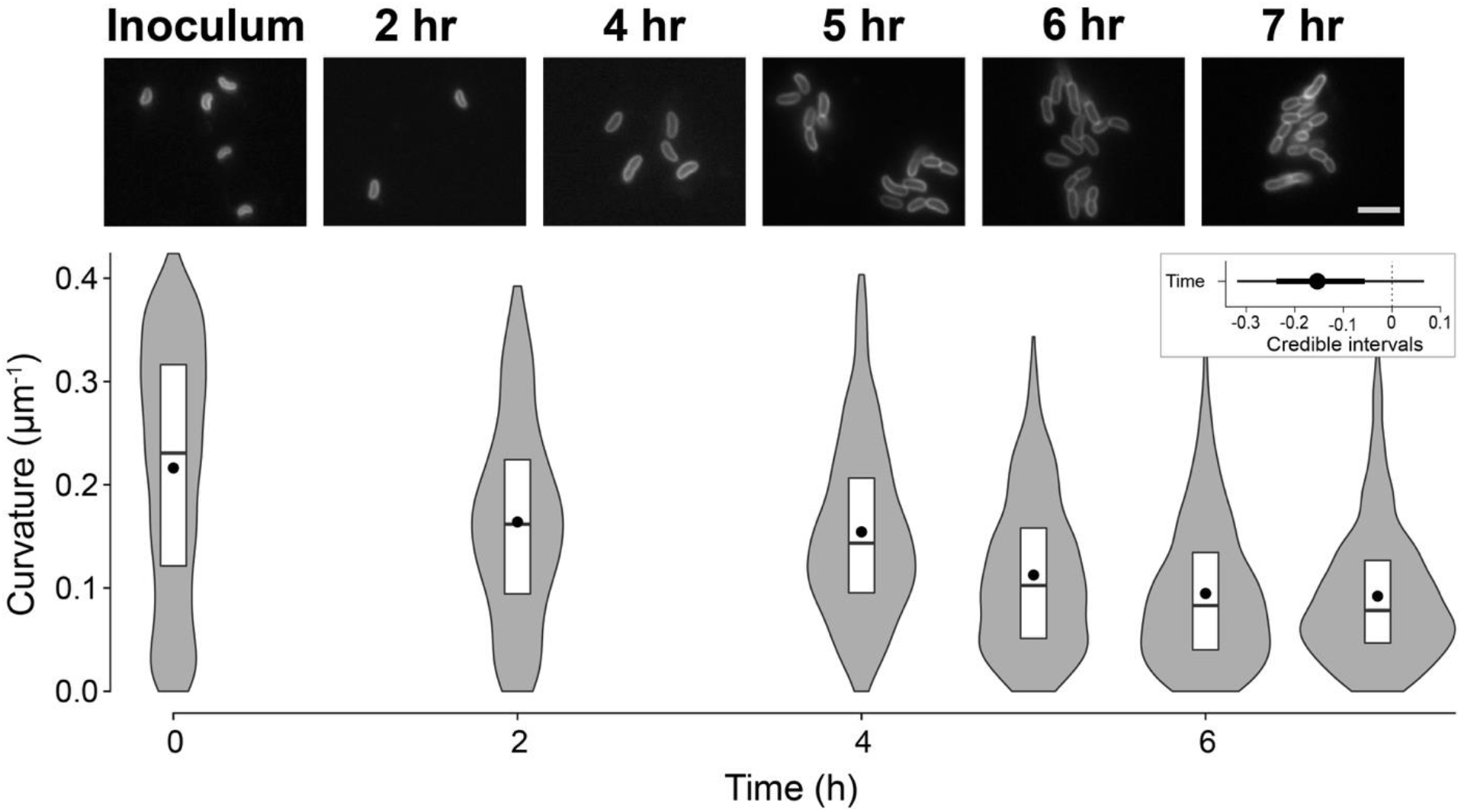
*V. cholerae* becomes straighter after attachment to a glass surface. **(A)** Representative images of *V. cholerae* from the inoculum and 2, 4, 5, 6, and 7 hours post attachment to glass coverslips. Cells were stained with FM4-64 prior to imaging. Scale bar is 5 μm. **(B)** Distributions of cell curvature at different time points after attachment. The box represents the first, second, and third quartiles. The dot represents the mean. Each distribution represents between 500 and 1,000 cells analyzed and pooled from two separate experiments. **Insert**) Credible intervals of the contribution of time to changes in the medians (dot = mean, thick line = 90% CI, thin line = 98% CI).

Our finding that *V. cholerae* transitioned from curved to straight rods during biofilm formation suggested this change in morphology is adaptive for biofilm formation. If so, we hypothesized that locking cells in a curved morphology, regardless of intracellular c-di-GMP concentrations, would negatively influence biofilm development. We tested this hypothesis by expressing CrvA from a multicopy plasmid using the *P*_*BAD*_ promoter in a *ΔcrvA* background. We found that basal expression of this construct (i.e. - no arabinose addition) was sufficient to restore curvature to the same level as wild-type planktonic cells and make cell shape insensitive to changes in c-di-GMP levels (Extended Data Figure 2). Thus, we generated three distinct populations: (1) cells that were able to transition normally between curved and straight rods (WT-pHERD20C), (2) cells that were constitutively straight (Δ*crvA*-pHERD20C), and (3) cells that were constitutively curved (Δ*crvA*-pCrvA) regardless of c-di-GMP concentrations (Figure 5A).

**Figure 5:**
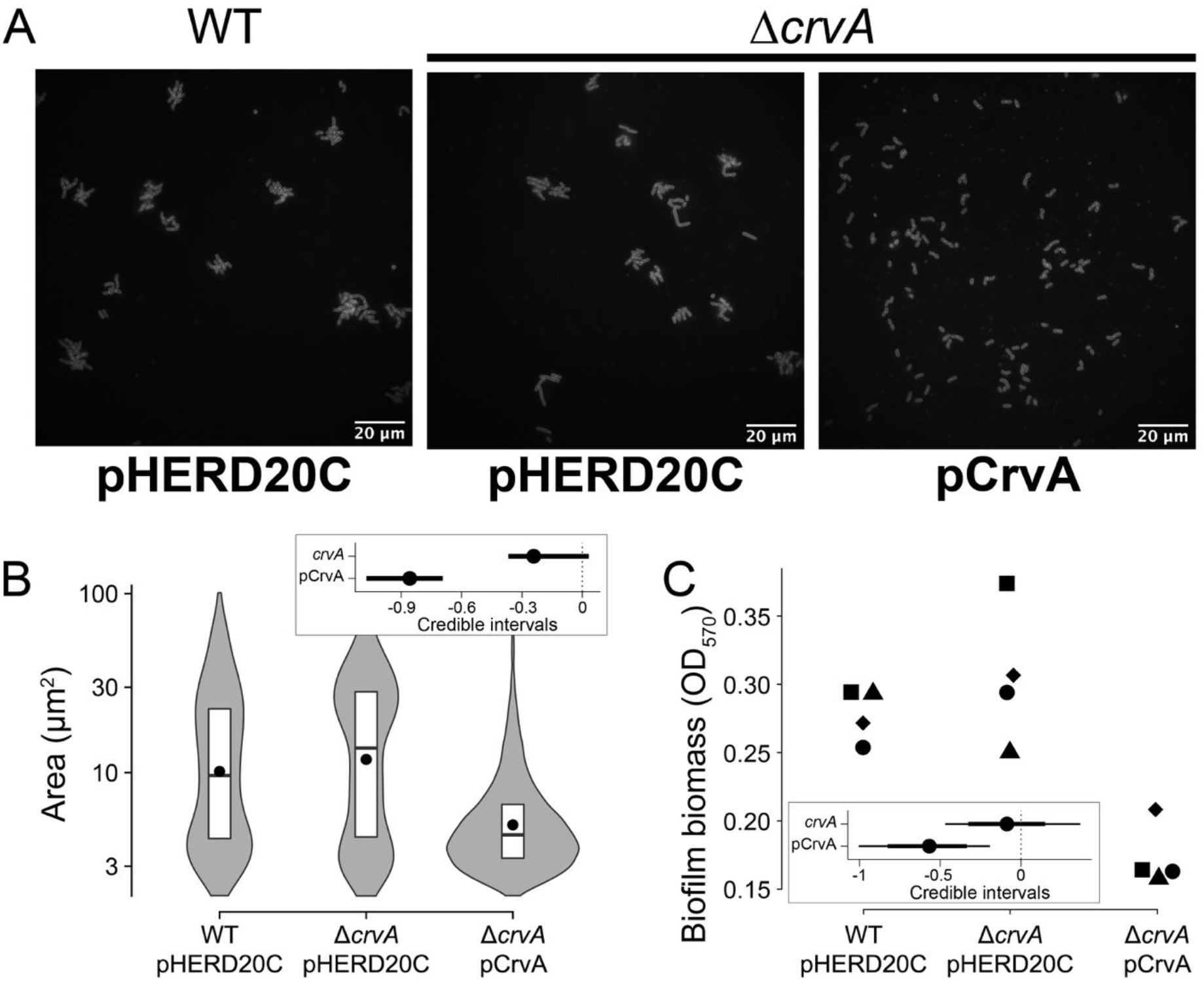
Curvature Influences Microcolony Development and Population Level Biofilm Formation. **A)** Representative images of *V. cholerae* microcolonies of WT harboring the control vector pHERD20C (left), *ΔcrvA* harboring the control vector pHERD20C (center), and *ΔcrvA* harboring the CrvA expression vector pCrvA (right). Microcolonies were stained with FM4-64 prior to imaging. Scales in the bottom right of each image are 20 μm. **B)** Distributions of microcolony areas pooled from two independent experiments totaling between 700 and 2000 microcolonies. The box represents the first, second, and third quartiles. The dot represents the mean. Each distribution represents between 500 and 1,000 cells analyzed and pooled from two separate experiments. **C)** Biofilm biomass formed by different strains. Dot shapes represent independent replicates. **Inserts**) Credible intervals of the contribution of *crvA* and pCrvA to changes in the medians (dot = mean, thick line = 90% CI, thin line = 98% CI).

We found that the area of individual microcolonies of the WT and the straight populations at 8 hours were 10 μm^2^ and 12 μm^2^ on average, indicating the straight mutant is not impaired in microcolony formation (Figure 5B). Microcolonies formed by the curved mutant were smaller than the size of WT (5 μm^2^, p-val < 2.5e-4) and contained only 2 to 3 cells (one cell = 2.6±1.4 μm^2^) (Figure 5B). To test if the difference in the areas of microcolonies could be accounted for by an inability to adhere, we quantified the total fluorescence emitted from cells attached on the glass surface for each mutant and observed that all strains had similar levels of surface coverage (Extended Data Figure 3).

**Extended Data Figure 3:**
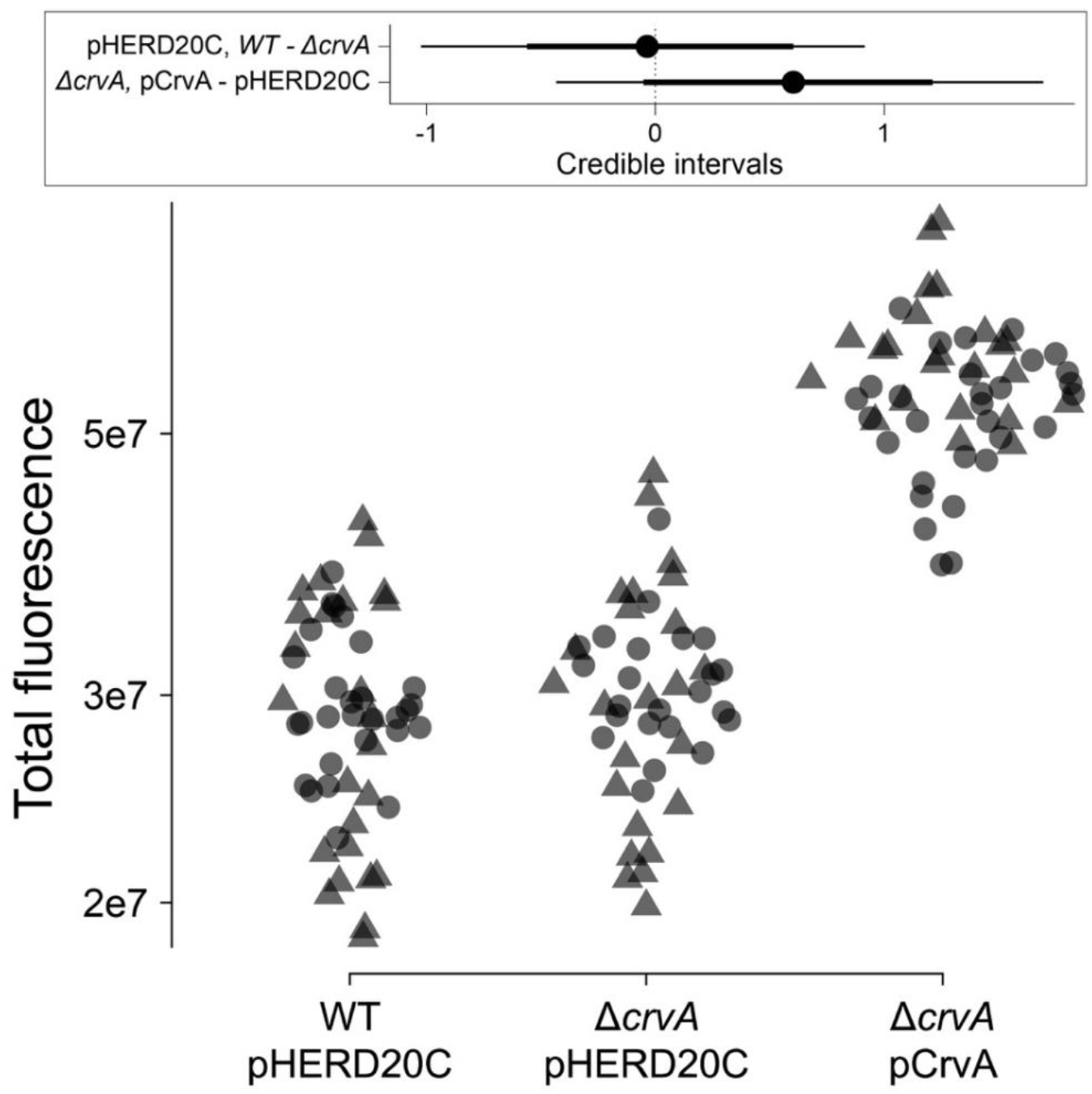
Effect of Curvature on Surface Coverage. Total fluorescence intensities from stained cells (FM4-64) attached to a glass surface for different strains. Each dot represents one field of view. Dot shapes represents independent replicates. **Insert**) Credible intervals of the differences in the means explained by *crvA* and pCRvA (dot = mean, thick line = 90% CI, thin line = 98% CI).

Based on the altered microcolony formation, we hypothesized that locking cells into curved rods would lead to a decrease in the biomass of mature biofilms. Using crystal violet to quantify the total biomass of attached cells from static culture in glass tubes, we found that WT and straight populations had indistinguishable biofilm biomass (Figure 5C) (difference in biomass, 95% CI = [−0.40, 0.22]). The curved mutant had reduced biofilm biomass compared to the other strains, supporting that reduced microcolony formation led to reduced overall biofilm biomass (Figure 5C) (difference in biomass, 95% CI = [−0.91, −0.28], p-val = 1.8e-3). Given that both of these strains adhered to the surface equal to the WT control and that lack of *crvA* causes no growth defects under similar conditions (7), we conclude that these differences in biofilm that we have observed are due to cell shape. Our analysis of cell curvature during biofilm formation indicates that cell shape is regulated processes and that decoupling of curvature from changes in c-di-GMP negatively affects biofilm formation in *V. cholerae*.

### Cell Curvature Increases the Swimming Speed of Flagellated Cells

Because wild-type *V. cholerae* is curved in growth conditions that promote a planktonic lifestyle, we hypothesized that curvature provides an advantage for cells swimming in liquid using flagellar motility. To test his hypothesis, we grew wild-type and the Δ*crvA* mutant in minimal medium supplemented with pyruvate to late exponential phase (most cells are highly motile in these conditions) and tracked single cells to quantify their swimming speed. We determined that curved rods swim on average 5.5% faster than straight rods (Figure 6A) (relative speed increase, 95% CI = [5.5%, 5.9%], p-val < 1e-5). The difference in the reversal frequency between the trajectories from each strain was not practically significant (relative difference in reversion frequency, 95% CI = [−0.87%, 2.8%]) (Figure 6B), indicating that cell curvature does not significantly affect cell behavior. Therefore, curvature is advantageous for motile cells by providing a significant boost in speed for bacteria that are already swimming relatively fast. Overall, our results support that *V. cholerae* modulates its cell shape by regulating CrvA expression through c-di-GMP signaling to increase its fitness both during biofilm formation and when swimming freely.

**Figure 6.**
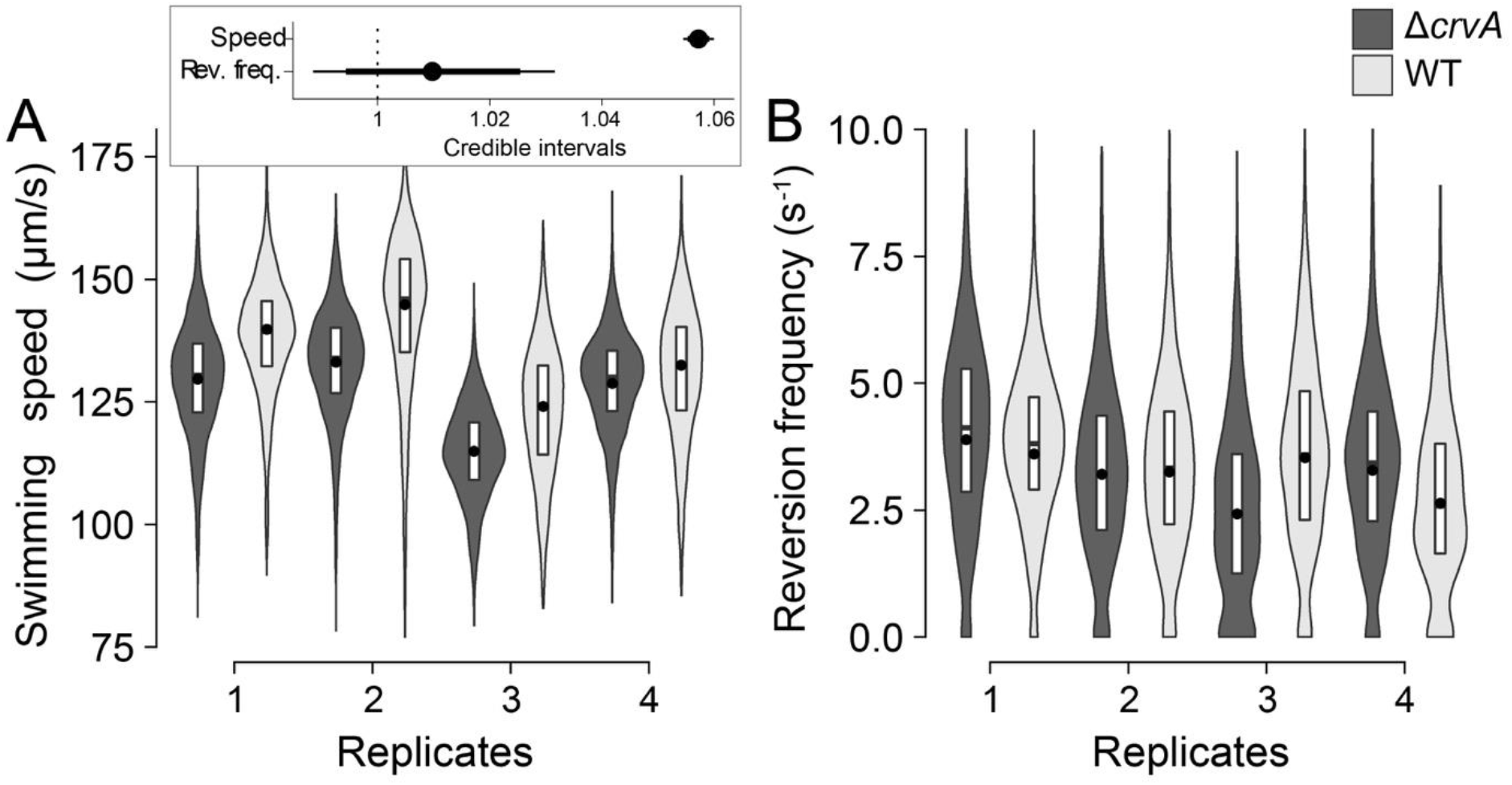
Curvature Increases the Swimming Speed of Flagellated Cells. Distributions of (A) swimming speed and (B) reversion frequency of the WT (light) and Δ*crvA* mutant (dark) determined by single-cell tracking on 4 independent replicates. The dot represents the mean. Each distribution represents more than 1,000 cell trajectories. **Insert**) Credible intervals of the relative difference in speed and reversion frequency between the two strains (dot = mean, thick line = 90% CI, thin line = 98% CI).

## Discussion

Derivatives of the rod shape, such as helical or vibrioid, are adaptations produced by genes with specific shape altering functions (2). While significant progress has been made understanding how bacteria build and manipulate their cell wall to alter cell shape, if and how these processes are influenced by environmental conditions is poorly understood. With multiple c-di-GMP synthesis and degradation enzymes that sense and responds to specific environmental cues, c-di-GMP signaling networks are one of the major mechanisms by which bacteria integrate environmental information to regulate their lifestyles, such as motility and biofilm formation. In this study, we show that c-di-GMP is a regulator of cell shape in the aquatic organism *V. cholerae*, where high c-di-GMP concentrations decrease cell curvature to generate straight rods. This response is dependent on the gene *crvA* and biofilm-promoting transcription factors, VpsR and VpsT, prompting us to assess the implications of shape in biofilm formation. We found that cells adopt straight morphologies while forming biofilms and that curvature retention caused defects in microcolony formation and mature biofilm production. Conversely, straight rods swim slower than curved cells when propelled by a single flagellum in liquid environments. This work highlights the importance of controlling cell shape when switching from a single-cell planktonic lifestyle to the development of multicellular communities such as biofilms. In addition, the regulation of *crvA* provides a novel example of cell-shape control by second messengers in bacteria.

Our conclusion that c-di-GMP negatively regulates curvature, *crvA* mRNA, and CrvA protein abundance is supported by several previous observations. *V. cholerae* utilizes quorum sensing (QS) as one mechanism to change intracellular c-di-GMP where cells in low-cell density (LCD) cultures have higher c-di-GMP concentrations than cells in high-cell density (HCD) cultures (26, 31). In 1928, the microbiology pioneer Alfred Henrici discovered that *V. cholerae* cell shape changes with culture density, stating: “The embryonic cells [those cells early in the growth curve] are therefore large, plump, and straight […] Therefore, the mature cells are slender and curved, the typical *Vibrio* form” (5). Additionally, Bartlett *et al* discovered curvature was a quorum regulated phenomenon (7). Studies comparing transcripts from cells in LCD to cells in HCD identified *crvA* as a QS regulated gene, with higher expression in the HCD state (32). Thus, the finding that cells at low density, which have high c-di-GMP concentrations, appear straight support our conclusion that c-di-GMP controls cell shape in *V. cholerae* (5, 7, 26).

The c-di-GMP-dependent transcription factor VpsT was sufficient to inhibit curvature in planktonic cells (Figure 2C), suggesting that VpsT is responsible for the straightening of cells during microcolony formation. While we could not directly test this, because *vpsT* is required microcolony formation, several studies strongly support this model (11, 21, 22, 27). Specifically, adhered cells produce VPS within 15 minutes and structural matrix proteins within an hour of surface attachment, both of which are positively regulated by VpsT and high c-di-GMP concentrations (28). Additionally, c-di-GMP production increases over time in surface attached *V. cholerae*, peaking around 6 hours after attachment (33). Similar increases in c-di-GMP have been observed for other bacteria such as *Pseudomonas aeruginosa* after surface attachment (34, 35). These studies support a model whereby upon attaching to a surface, c-di-GMP concentrations increase and activate the VpsR/VspT regulatory network, resulting in surface adhesion and changes in cell curvature during biofilm development.

Pandemic isolates of *V. cholerae* often retain their vibrioid morphology when examined microscopically (36, 37). However, Wucher and co-workers found that nutrient deprivation in pandemic isolates induced cell filamentation, which aided in chitin colonization and VPS-independent biofilm formation under flow conditions (37). While the cells were longer than the reference strain, they appeared to retain their curvature, suggesting this mechanism is independent of *crvA* and c-di-GMP (7, 37). Single *V. cholerae* cells in VPS-dependent microcolonies, however, appear straight rather than curved in submerged flow cell conditions in multiple reports, yet the degree of change in cell shape was never quantified (28–30). Thus, our work is the first to quantify curvature in two-dimensional developing microcolonies to demonstrate that surface adhered cells become straighter as the biofilm develops (Figure 5).

While *V. cholerae* curvature has been shown to promote motility in plate-based motility assays, how shape influences specific parameters of motility such as chemotaxis or swimming speed at the single-cell level was not determined (7). Our findings indicate the influence of cell shape on motility extends beyond promoting movement in restricted spaces such as the matrices formed in low-agar motility assays. By tracking single cells swimming in liquid media, we found that constitutively straight cells swim slower than curved cells (Figure 6). Our study provides experimental evidence supporting theoretical research that predicted curvature improves cellular swimming efficiency in solution by optimizing rotational and translational resistance, resulting in more power transferred to the flagellum (38).

Examples of vibriod or helical morphologies include *C. crescentus, Helicobacter pylori,* and *V. cholerae*, and many vibrioid shaped bacteria are found in aquatic reservoirs (39). Curvature is likely to influence how each organism adapts to their respective niche. For example, curvature in *C. crescentus* is important for positioning the cell pole near the surface during biofilm growth in flow conditions as straight mutants are unable to form biofilms under these conditions, which is opposite of the conclusions for *V. cholerae* drawn here (40). The helical shape of the human pathogen *H. pylori* is thought to be an adaptive trait that promotes fitness in a human host, however straight *H. pylori* clinical isolates have been identified that provide distinct advantages and disadvantages in certain physiological niches during infection (41). These results emphasize that cell shape can differentially impact bacterial species based on their own specific physiologies

Rod-shaped cells are observed in all three domains of life, suggesting this shape provides evolutionary advantages compared to others (42). For example, mutations that alter the morphology of the rod-shaped bacteria *E. coli* or *Rhodobacter sphaeroides* also negatively impact biofilm formation, which could impact fitness in certain environments (43). Further, analysis of cell packaging in dense clusters highlighted the importance of the rod shape in cell positioning and self-organization in developing microbial communities (44, 45). While helical or curved bacteria like *H. pylori* and *C. crescentus* can access the adaptive benefits of curved or straight cells through mutagenesis and loss of function, *V. cholerae* has evolved to accomplish this by utilizing environmental sensing and c-di-GMP signaling (40, 41). Our work suggests cell shape in bacteria is a malleable feature that is regulated by environmental signaling to maximize swimming speed in the planktonic state and cell-surface interactions during biofilm development.

## Materials and Methods

### DNA manipulations and growth conditions

*V. cholerae* C6706 Str2 was used in all experiments. In Figures 1 through 3, the biofilm mutant derivative (*ΔvpsL*) was used as the parental strain to prevent aggregate formation at high c-di-GMP so that individual bacteria could be imaged in solution. The *ΔvpsT* and *ΔvpsR* mutants as well as expression vectors for VpsT and VpsR were previously generated (12). pKAS32 was used to construct deletion and knock-in strains (46). Unless otherwise stated, all cloning was done by Gibson Assembly (NEB). For the deletion construct pKAS32_ΔcrvA, pKAS32 was digested with XbaI and SacI and purified by gel extraction (Promega). Primers (Integrated DNA Technologies) were designed with NEBuilder (www.nebuilder.com, NEB) to incorporate the appropriate 5’ ends necessary for Gibson Assembly and 3’ gene specific ends. Specifically, 700 bp upstream and downstream of *crvA* (VCA1075) were amplified by PCR (Q5 Polymerase, NEB) using *V. cholerae* gDNA as a template. For the knock-in construct pKAS32_crvA-HIS, the replacement allele harboring a 6-histidine tag before the stop codon was amplified by PCR with the HIS tag incorporated into the 5’ of the oligonucleotide primer. For the knock-in construct pKAS32_pcrvA-CrvA, the native promoter and ORF of *crvA* (−358 bp relative to the ATG start codon until the translational stop codon) were amplified using PCR and inserted into the locus VC1807, a pseudogene encoding an authentic frameshift that provides a neutral site for insertion into the genome (47). For pBBRLux_pcrvA, the upstream sequence of *crvA* from −358 to −1 bp relative to the ATG start codon was amplified and Gibson cloned into the BamHI and SpeI restriction site of pBBRlux. For pHERD20C_CrvA, the ORF of *crvA* was amplified and cloned into the KpnI and SacI sites of pHERD20_C. Successful clones were screened by colony PCR using GoTaq polymerase (Promega) and sequenced by Sanger sequencing (Genewiz Inc.) to ensure no mutations were incorporated during the cloning process. Construction of knock out and knock in strains were carried out by the protocol for Skorupski *et al*. (46). Plasmids were moved from S17-λpir *E. coli* into *V. cholerae* by conjugation using Polymixin B for counterselection (10 U/mL). Unless otherwise stated, both *E. coli* and *V. cholerae* were propagated in LB with ampicillin (100 μg/mL), kanamycin (100 μg/mL), and/or chloramphenicol (10 μg/mL) when needed. The inducer isopropyl-β-D-thiogalactoside (IPTG) was routinely used at 100 μM unless stated otherwise.

### Phase contrast microscopy and single-cell analysis

Overnight cultures of *V. cholerae* were diluted 1:100 into 2 mL LB supplemented with ampicillin and IPTG at appropriate concentrations. Cultures were grown until an OD_600_ of 1.3 to 1.5, at which point cells were diluted to an OD_600_ of 0.5 in microcentrifuge tubes. 1% agarose pads in deionized water were cut into squares of approximately 20 × 20 mm and placed on microscope slides (75 × 25 × 1.0 mm (L×H×W), Alkali Scientific Inc.). 2 μL of diluted cultures were spotted onto glass coverslips (22 × 22 mm, #1.0 thickness, Alkali Scientific) and the coverslip was gently placed onto the agarose pad. Phase-contrast microscopy was carried out with a Nikon Eclipse Ti-E inverted microscope equipped with a 100X phase contrast oil immersion objective (1.4 NA), a Nikon Perfect Focus System, a Prior H117 ProScan motorized stage, a Lumencor PEKA white LED diascopic light source, and an Andor Zyla 4.2 sCMOS camera. The microscope and camera were controlled by a computer workstation with MATLAB (Mathworks Inc.) and Micromanager version 1.4 (micro-manager.org). Uneven illumination and image artifacts were corrected using a flat field image of a clean slide. Cells were detected and segmented using the Fiji plugin MicrobeJ with the following settings: area (0 – 4.5, μm^2^), length (1.5 – max, μm) and width (0 – 3, μm) (48). Data from segmented images were analyzed using R to plot curvature (μm^−1^), width (μm), and length (μm). Representative images were cropped, and the scale bar was added using Fiji software (49). The median cell length, width, and curvature was calculated for each biological replicate. The posterior probability distributions and the p-values for the different factors tested (QrgB, VpsR, VpsT, and CrvA inductions) were calculated for the medians using different linear mixed-effect models with a lognormal distribution link function. Figure 1 and ED1: (Curvature, Length, Width) ~ QrgB * IPTG + (QrgB * IPTG|Replicate), Figure 2: (Curvature, Length, Width) ~ Complement + QrgB: Complement + (Complement + QrgB: Complement |Replicate), Figure 4: (Curvature, Length, Width) ~ Time + (Time|Replicate).

#### Analysis of curvature of attached cells in microcolonies

Overnight cultures were diluted 1:1000 in 1 mL 1X phosphate buffered saline (PBS, Sigma) by 10-fold serial dilutions. UV-sterilized #1 cover slips (22×22 cm) were placed in 6-well plates (Costar®) submerged in 1 mL LB. Cultures, in six sets of biological replicates, were seeded in the wells with slides by further diluting 5-fold, resulting in a final dilution of 1:5000, followed by gentle swirling. Microcolonies were allowed to develop during static incubation at 21°C. At the given timepoint, the media for two biological replicates were removed by aspiration, the wells washed with 1 mL 1X PBS, and the resulting adhered bacteria on the cover slip were stained with 200 μL of the membrane stain N-(3-Triethylammoniumpropyl)-4-(6-(4-(Diethylamino) Phenyl) Hexatrienyl) Pyridinium Dibromide (FM4-64) (Sigma) at a final concentration of 20 μg/mL for 5 minutes. The stain was removed by an additional wash with 1 mL 1X PBS. Small sections of agarose pads (~5×5 mm) were arranged in a 20×20 cm square pattern on glass microscope slides. The glass coverslip containing the stained microcolonies was then inverted and placed on top of the agarose pads. Biofilm imaging was carried out using a Leica DM5000b epifluorescence microscope with a 100X-brightfield objective (1.4 NA) equipped with a Spot Pursuit CCD camera and an X-cite 120 Illumination system. Images were acquired using the dsRed filter set (Excitation 560/40 nm, Emission 620/60 nm). Each slide was imaged with at least 20 fields of view for each biological replicate at each timepoint. Cells within microcolonies were manually outlined in MicrobeJ using the manual interface option with at least 500-1000 cells outlined per replicate. Data from MicrobeJ analysis were exported into R for analysis. The posterior probability distribution of the effect of time on the morphology of attached cells was calculated using a linear mixed-effect model with a lognormal distribution link function ((Curvature, Length, Width) ~ Time + (Time|Replicate)).

#### Microcolony Area Analysis

For single timepoint biofilm analysis, 8-well microchamber slides (μ-Slide, 8-well glass bottom, ibidi) were used. 1:1000 diluted overnight cultures were additionally diluted 1:5 in an individual well of the microchamber slide in 200 μL LB supplemented with chloramphenicol at the appropriate concentration. Each slide had three strains in biological duplicate. Microcolonies were developed by incubating the microchamber slide statically at 21°C for 8 hours, resulting in WT microcolony sizes between 10-20 μm^2^. Media was removed from slides by aspiration, each well was washed with 200 μL 1X PBS, and the microcolonies were stained with FM4-64 (150 μL) at a final concentration of 20 μg/mL for 5 minutes. The remaining stain was washed with 200 μL 1X PBS and biofilms were imaged by inverting the microchamber (positioning the glass bottom upwards) by fluorescence microscopy using a Leica DM5000b epifluorescence microscope as previously described. At least 20 fields of vision were captured per replicate per strain and the resulting images were processed using Fiji (Enhance Contrast, Saturated Pixel % - 0.3 (49)). Processed images were then analyzed using MicrobeJ with the following settings: Background type (Dark), Mode of detection (Basic), Area (4.5 – 100, μm^2^), and circularity (0-1). Data were exported into R for analysis. For all fluorescence microscopy, representative images were cropped, and the scale bar was added using Fiji software (49). The posterior probability distributions of the effects of mutation and complementation of the microcolonies areas were calculated using a linear mixed-effect model with a lognormal distribution link function (Area ~ Strain + Complement + (Strain + Complement |Replicate)). For measurement of total surface fluorescence, images above were processed using Fiji (Subtract Background, Rolling ball radius: 50 pixels (49)) and analyzed for total integrated density using the Measurement function in Fiji. Data were exported into R for analysis.

### Luciferase Reporter Assay

Overnight cultures of four biological replicates of *V. cholerae* harboring *P*_*crvA^−^*_ transcriptional fusions to luciferase in pBBRlux were diluted 1:100 in 1 mL LB supplemented with ampicillin, chloramphenicol, and IPTG in 1.5 mL microcentrifuge tubes. 150 μL of cell solution was aliquoted into wells of a black 96-well plates (Costar) in technical duplicates. Plates were incubated at 35°C with shaking at 220 RPM. Every hour, luciferase (Relative Light Units) and OD_595_ measurements were taken, and luciferase activity was measured using an Envision plate reader (Perkin Elmer). The posterior probability distributions of the relationship between the light emission and optical density was calculated using a linear mixed-effect model with a normal distribution link function (RLU ~ OD_600_:QrgB + (OD_600_:QrgB|Replicate)).

### RNA Isolation and qRT-PCR

RNA isolation and qRT-PCR analysis were carried out following protocols described previously (20). In brief, overnight cultures were diluted to a starting OD_600_ of 0.040 in 2 mL LB supplemented with ampicillin and IPTG and grown until an OD_600_ of approximately 1.3 at 35°C and 220 RPM. 1 mL of each replicate was pelleted, and RNA was extracted using the TRIzol^®^ reagent following the manufacturer’s directions (Thermo Fischer Scientific). Purified RNA was quantified using a NanoDrop spectrophotometer (Thermo Fischer Scientific). 5 μg of purified RNA was treated with DNAse (Turbo DNAse, Thermo Fischer Scientific). cDNA synthesis was carried out using the GoScript™ Reverse Transcription kit (Promega). cDNA was diluted 1:30 into molecular biology grade water and amplification was quantified using SYBR Green (Applied Biosystems™). Reactions consisted of 5 μL 2.5 μM primer 1, 5 μL of 2.5 μM primer 2, 5 μL of diluted cDNA template, and 15 μL of 2X SYBR green consisting of dNTPs and AmpliTaq Gold® DNA polymerase. Each plate had technical duplicates and biological triplicate samples as well as no reverse transcriptase reactions to detect genomic DNA contamination. The StepOnePlus Real Time PCR system was used for qRT-PCR with the following thermocycling conditions: 95°C for 20 seconds then 40 cycles of 95°C for 2 seconds and 60°C for 30 seconds. Melting curves were included to ensure PCR products had single amplicons. Data was analyzed by the ΔΔCt method using *gyrA* as a reference target. The experiment was repeated with 10 biological replicates on separate days and the data from each experiment were pooled.

### Western blot analysis

Overnight cultures of the parent or *ΔvpsT* backgrounds with 6-histidine tagged *crvA* integrated at the *crvA* locus (CrvA-HIS) harboring QrgB* or QrgB were diluted 1:100 in 10 mLs LB supplemented with ampicillin and IPTG in 50 mL baffeled flasks. Cultures were grown until OD_600_ reached ~ 1.3, collected by centrifugation (2.5 minutes at 7,000 x g), resuspened in 200 μL lysis buffer (20 mM Tris-HCl, 1% SDS, pH 6.8), and moved to 1.5 mL microcentrifuge tubes, and boiled for 10 minutes at 95°C. Boiled lysates were centrifuged (1 minute at 12,000xg) to pellet insoluble material and the supernatants were moved to new 1.5 mL microcentrifuge tubes. Samples were diluted 1:20 in 1X PBS and protein concentration was quantified by the DC protein assay with BSA as the standard (BioRad®). Lysates were normalized to 5 μg/μL in 2X loading buffer (Lysis buffer supplemented with 5 mM beta-mercaptoethanol and Coomassie blue). 20 μL of normalized lysates were loaded into pre-cast 4-20% SDS-PAGE gels (4-20% Mini-PROTEAN^®^ TGX™ Precast Protein Gels, BioRad) alongside size standards (Precision Protein Plus™, BioRad). Gels were run at room temperature for 90 minutes at 90 volts in 1X Tris/Glycine/SDS running buffer (BioRad). Blots were transferred to nitrocellulose membranes using the Mini Trans-Blot® system (BioRad) with TBST (Tris buffered saline with Tween 20, pH 8.0)-methanol (20% v/v). Transfers were carried out at 4°C for 2 hours at 250 mAmps. After the transfer, blots were removed and placed in blocking buffer (TBST supplemented with 5% skim milk) and incubated with agitation for 1.5 hours at room temperature. Blocking buffer was removed and replaced with 20 mL blocking buffer supplemented with 4 μL anti-HIS primary antibody (mAb, mouse, GenScript®) and the blot was incubated overnight at 4°C with agitation. The next day, the blot was washed three times by removing spent blocking buffer, addition of 20 mL blocking buffer, and agitation for 2-3 minutes at room temperature. After the last wash, 20 mL blocking buffer supplemented with 5 μL of horseradish peroxidase (HRP) conjugated rabbit anti-mouse secondary antibody (BioRad) was added and incubated for 2 hours at room temperature with agitation. After incubation with the secondary antibody, the blot was washed with 5 cycles of 10 mLs blocking buffer with 1-minute agitations. Reagent 1 and 2 from the Pierce ECL kit were mixed according to manufacturer’s instructions and chemiluminescence was detected using Amersham Imager 600 with the ‘Chemilumescence’ settings. Images were removed from the imager, uploaded to Fiji, cropped to remove the protein standards, and were processed by enhancing contrast (Enhance Contrast, 0.3% saturated pixels). Band intensities were measured using the Gel Analyzer function. Two separate experiments with two biological replicates were repeated on separate days (n = 4). The posterior probability distributions of the effects of VpsT and QrgB on the accumulation of CrvA-His were calculated using a linear mixed-effect model with a lognormal distribution link function (Intensity ~ QrgB*VpsT + (QrgB*VpsT|Replicate)).

#### Crystal Violet Biofilm Assay

Overnight cultures of *V. cholerae* were diluted 1:100 into 1 mL LB supplemented with chloramphenicol into new, sterile 18 × 150 mm borosilicate test tubes and statically incubated at 21°C for 8 hours. After 8 hours, media and unattached bacteria were removed, and the biofilms were washed twice with 1 mL 1X PBS (PBS washes were removed by aspiration). 1 mL crystal violet (CV) (0.4%) solution was added to each tube and the biofilm was stained for 10 minutes. The stain was removed by aspiration and the stained biofilm was washed twice with 1 mL 1X PBS. CV was eluted with ethanol and the absorbance at 570 nm (OD_570_) was measured. The posterior probability distributions of the effects of mutation and complementation of the biofilm biomass were calculated using a linear mixed-effect model with a lognormal distribution link function (Biomass ~ Strain + Complement + (Strain + Complement |Replicate)).

#### Tracking and analysis of swimming cells

Wild-type and *crvA* mutant cells were grown in M9 minimal salts (52 mM Na_2_HPO_4_, 18 mM K_2_HPO_4_, 18.69 mM NH_4_Cl, 2 mM MgSO_4_, pH 7) supplemented with 10 μM FeSO_4_, 20 μM C_6_H_9_Na_3_O_9_ and 36.4 mM Sodium pyruvate shaking (200 r.p.m.) in liquid cultures at 37°C. Cultures were sampled at early stationary phase (1.9 × 10^9^ c.f.u./mL) and diluted in fresh medium to 10^7^ c.f.u./mL and incubated for 15 minutes before tracking. Swimming cells were tracked in liquid following the protocol previously described (50). Briefly, polyvinylpyrrolidone (PVP) was added at 0.05% w/v to the samples to prevent attachment on the glass slide. 6 μl of each sample were dropped on a glass slide and trapped under a 22 × 22 mm, #1.5 coverslip sealed with wax and paraffin to create a thin water film (10±2 μm) for video microscopy. The samples were kept at 37°C during tracking. Images of swimming cells were recorded using a sCMOS camera (Andor Zyla 4.2, Oxford Instruments) at 20 frames per second using a 40X objective (Plan Fluor 40x, Nikon Instruments, Inc.) mounted on an inverted microscope (Eclipse Ti-E, Nikon Instruments, Inc.). Cell were illuminated using phase contrast. Images were analyzed to detect and localize cells using custom scripts (50) and cell trajectories were reconstructed using the μ-track package (51). The analysis of the cell trajectories was done in MATLAB (The Mathworks, Inc.) as previously described (50). The mean swimming speed of each cell was calculated by averaging instantaneous speed along the trajectory when the cell is not doing a reversal. The posterior probability distributions and the p-vals for the differences in mean swimming speed and reversal frequency for each strain were calculated using a linear mixed-effect model with a lognormal distribution link function ((Mean speed, Rev. freq.) ~ Strain + (1|Replicate)).

#### Statistical analyses

All the statistical analyses were done using Bayesian sampling of the respective mixed-effect generalized linear models using the RSTAN (52) and BRMS packages (53) in R (54) with 4 chains, each with 1,000 warmup iterations and at least 2,500 sampling iterations. Uninformative priors were set to the defaults generated by BRMS. The plots were generated using the ggplot2 and tidybayes packages (55, 56).

## Data availability

All primary data supporting the results described in this study are available from the corresponding upon reasonable request.

## Acknowledgments

This work was supported by NIH grants GM109259, GM110444, and AI143098 to C.M.W., NSF grant 1714612 to Y.S.D and C.M.W., and funds from Michigan State University to Y.S.D.

## Strains, Plasmids, and Primers

**Table 1:**
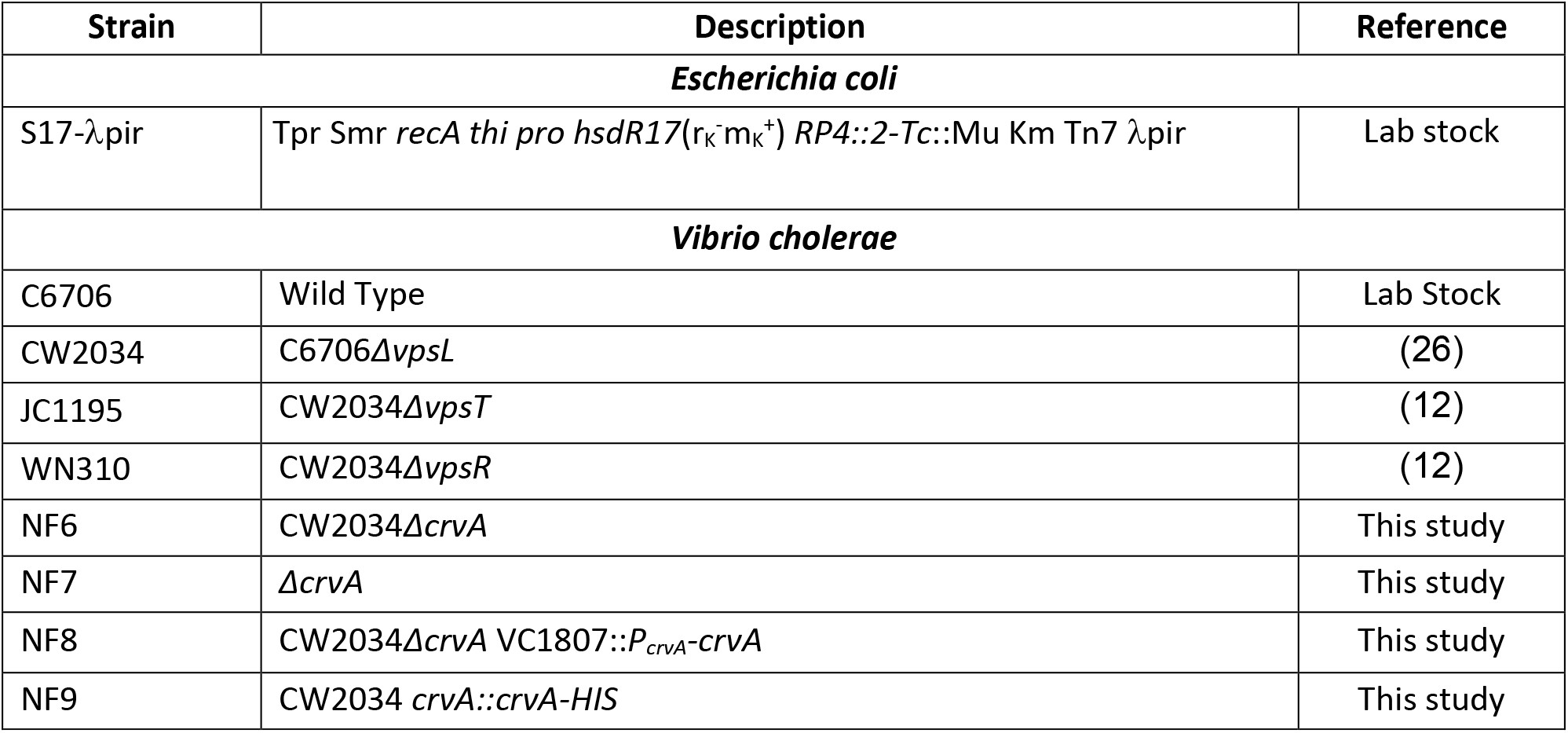
Strains Used in this Study.

**Table 2:** Plasmids Used in this Study.

| Plasmid | Description | Reference |
| --- | --- | --- |
| pBBRlux | *luxABCDE* containing promoter-less plasmid, Cam<sup>R</sup> | Lab collection |
| pEVS143 | *pTac* overexpression vector, Kan<sup>R</sup> | (57) |
| pKAS32 | Sucide vector for mutant construction, Amp<sup>R</sup> | (46) |
| pBRP333 | Control expression vector, Kan<sup>R</sup> | (18) |
| pBRP1 | *qrgB* * (inactive DGC) cloned into expression vector pMMB67Eh, Amp<sup>R</sup> | (18) |
| pBRP2 | *qrgB* (active DGC) cloned into expression vector, pMMB67Eh, Amp<sup>R</sup> | (18) |
| pCMW131 | *vpsR* in pEVS143, Kan<sup>R</sup> | (19) |
| pCMW132 | *vpsT* in pEVS143, Kan<sup>R</sup> | (19) |
| pHERD20T | *P<sub>BAD</sub>* Expression vector | (58) |
| pNF057 | pHERD20T *pBAD* expression vector with Cam<sup>R</sup> replacing Amp<sup>R</sup> | (20) |
| pNF069 | 358 bp upstream of ATG of *crvA* (*P<sub>crvA</sub>*) in pBBRlux | This study |
| pNF070 | Deletion construct for *crvA* in pKAS32, Amp<sup>R</sup> | This study |
| pNF071 | VC1807 knock-in construct, Amp<sup>R</sup> | This study |
| pNF072 | VC1807 knock-In construct with CrvA driven by native promoter *P<sub>crvA</sub>*, Amp<sup>R</sup> | This study |
| pNF073 (pCrvA) | CrvA from C6706 gDNA into pNF057 | This study |

**Table 3:**
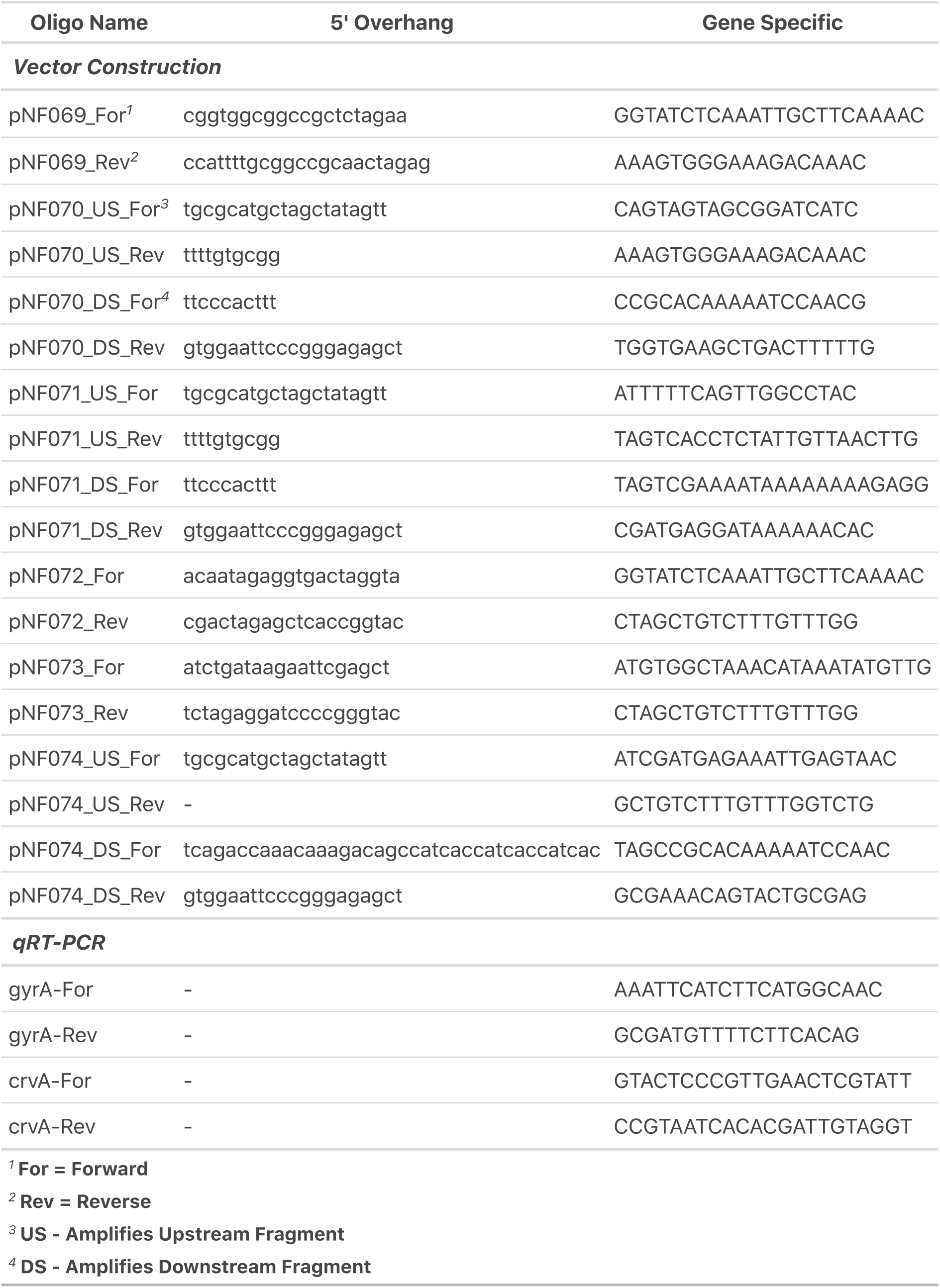
Oligonucleotides Used in this Study.

